# Varia: Prediction, analysis and visualisation of variable genes

**DOI:** 10.1101/2020.12.15.422815

**Authors:** Gavin Mackenzie, Rasmus W. Jensen, Thomas Lavstsen, Thomas D. Otto

## Abstract

Assessing the diversity or expression of variable gene families in pathogens can inform about immune escape mechanisms or host interaction phenotypes of clinical relevance. However, obtaining the sequences and quantifying their expression is a challenge. Here, we present a tool, which based on unique sequence tag similarity between members of a gene family, predicts the domains encoded by the queried gene. As an example, we are using the *var* gene family, encoding the major virulence proteins (PfEMP1) of the human malaria parasite, *Plasmodium falciparum*. We developed Varia, which predicts the likely *var* gene sequence and encoded protein domain composition of a gene from short sequence tags. We provide a new extended annotated *var* genome database, in which Varia identifies genes with identical tag sequences and compares these to return the most probable domain composition of the query gene. Varia’s ability to predict correct PfEMP1 domain compositions from short *var* sequence tags was tested in two complementary pipelines to (a) return the putative gene sequences and domain compositions of the query gene from any partial sequence provided, thereby enabling detailed assessment of specific genes’ putative function and experimental validation of these (b) to accommodate rapid profiling of *var* gene expression in complex patient samples, by compiling the overall domain prevalence among *var* transcripts predicted identified and quantified by next generation sequencing of so-called *var* DBLα-sequence tags.

**Availability and implementation:** Varia is available on GitHub (https://github.com/GCJMacken-zie/Varia) under the MIT license.

## 1 Introduction

Pathogens can evade the immune system through polymorphic protein families interacting with host molecules. In *Plasmodium falciparum* malaria parasites, the *var* genes encode the *P. falciparum* membrane protein 1 (PfEMP1) family, which members are inserted into the erythrocyte membrane to bind select human endothelial receptors and enable the parasites escape from blood circulation and splenic destruction. The extra-cellular part of PfEMP1 are comprised multiple DBL and CIDR domains (supplementary data) and separate receptor-binding phenotypes and clinical outcomes of infection have been associated with specific subtypes of DBL and CIDR domains^1^. PfEMP1 are important targets of acquired immunity to malaria^2^, and hence the PfEMP1 protein family have diversified to escape immune recognition. Each parasite encodes 50-60 *var* genes conferring the parasite a similar repertoire of receptor-binding phenotypes Rask ^3^. Recently, through whole genome sequencing of 2400 parasites collected across the world, a database of over 140,000 *var* genes was generated^4^. Here, we sought to exploit this unprecedented sequence depth, and extend it with further 750 samples to build a tool, which will enable to reconstruction and experimental validation of the near full-length *var* genes from any sequence tag available; and which can provide rapid quantification of PfEMP1 domain-specific expression in complex malaria patient samples by analysis of *var* DBLα expression tags.

## 2 Methods

Varia is programmed in Bash and Python and results are visualized using Circos and Excel readable files. The tools offers two analysis pipelines. In the *Var* Identification and Prediction (Varia_VIP) pipeline, the user provides one or more partial *var* sequence(s), which are used for searching a comprehensive *var* database for near identical sequences. Hit sequences are clustered based their full-length sequence similarity, and the domain composition of the longest gene sequence representing each cluster is visualized in circular plots (Figure 1A) and tabular output files (see supplementary data).

**Figure 1.**
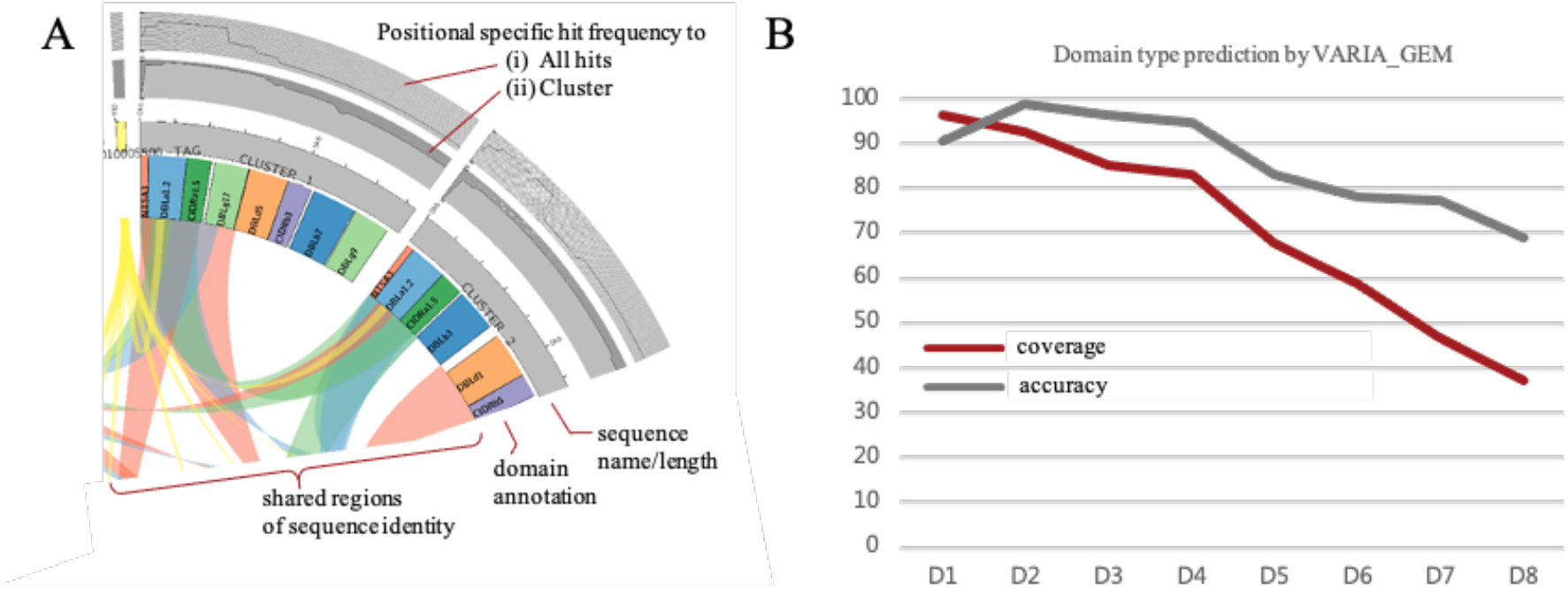
Examples of Vaira output and performance. A) Section of a Circos plot from Varia_VIP showing sequence similarities and domain traits of database genes similar to a query DBLα tag (yellow). B) Graphs showing accuracy (correctly predicted domain types) and coverage (proportion of query DBLα tags for which predictions could be mad) per PfEMP1 domain position D1-D10.

In the Varia Gene Expression Module (Varia_GEM), batches of DBLα-tag sequences e.g. from as high-throughput sequencing data from multiple patient samples are clustered to show the distribution of unique DBLα-tag sequences in each data batch. For each unique DBLα-tag, near identical sequences in the *var* database are identified and their domain composition processed to generate a consensus prediction of the domain composition of each query gene and the relative expression level of all known PfEMP1 domains in each batch of sequences. The output is given in Excel file format allowing subsequent statistical analyses of domain type association with e.g. clinical data pertaining to the samples (see supplementary data).

For Varia a new *var* genome database containing all assembled *var* gene contigs including their domain annotations from 2400 Illumina sequenced and 15 Pacbio sequenced^5^ as well as additional newly assembled var genes of further 755 isolates was build and provided with the tool.

## 3 Results

The tool’s ability to predict *var* gene domain annotations was first tested using 150 bp sequence tags derived from DBL domains of *var* genes from 15 curated and annotated *P. falciparum* genomes using the Varia_VIP pipeline. As Varia_VIP returns all possible sequence and domain compositions of the query gene based on database sequences with near identical tags, the number of different gene predictions made, was assessed at different similarity thresholds for the database search (Table S1). As expected, lowering the similarity threshold increased the proportion of queries with correctly predicted annotations but at the cost of a higher number of different predictions. Specifically, for the DBLα domain tags, which are found in the 5’end all *var* genes, a correct annotation was identified for 72% of the query genes at 99% sequence similarity threshold. The correct annotation was found among on average 14 suggested predictions. At this 99% similarity threshold, prediction of the exact query DNA sequence, defined as 99% identity over at least 80% of the sequence, was successful for 29% of the DBLα domain tags. Lowering the similarity threshold to 90% resulted in a limited increase to 78% of the query genes with correct annotation among the suggested domain compositions, and an increase in the number of alternative suggestions. For this reason, we recommend applying the highest similarity threshold resulting in database hits when using Varia_VIP to search for putative domain compositions and sequences of query genes, to allow manageable experimental validation (e.g. by PCR) of the query sequence.

In some studies, for example of *var* gene expression in patients, a rapid prediction of a single PfEMP1 domain composition from a large number of sequences, is required. For this, we build Varia_GEM, which based on all hit sequences to a DBLα-tag returns the likely consensus domain annotation of the query gene (supplementary data 4). In short, for each domain position (D1-D10; DBLα is always at position D2), the tool determines if a specific PfEMP1 main domain type or domain subtype is dominant (>66%) among hit genes sequences. If this is the case, the tool returns the consensus annotation in a tabular format along with quantitative data of frequency of sequence tags associated with the domain or domain composition to allow quantitative analysis of PfEMP1 domain traits with e.g. clinical features pertaining to the origin of the sequences. Based on testing the accuracy and rate (coverage) of predictions at different parameter settings, a sequence similarity threshold of 95% over 200 base pairs was chosen as default (Figure 1B). This setting results in >90% of domains at position D1-D4 and 70-80% of domains at D5-D8 position being correctly predicted, and an ability to predict domains (coverage) at position D1-D4 in >80% of queries. Prediction rates dropped markedly from position D5-D10.

## 4 Discussion

Due to the diversity of PfEMP1 sequences, resolution of unknown full-length *var* gene sequences and their domain structure has required cumbersome laboratory proceedings. With Varia, we generated a tool that predict *var* gene sequences from small easily obtained sequence fragments, to allow the community to reconstruct genes of interest. The likelihood of correct predictions of sequence and domains flanking the query tags was high, whereas predictions distal to the query tag was less likely. These limitations are caused by the sequence diversity and recombination history of the *var* genes. With awareness to these limitations however, sequences and domain compositions of *var* genes of interest can be experimentally validated by PCR and differential *var* type expression between groups of patient samples can be assessed. The Varia tool will thus be useful to understand the distribution and clinical importance of different *var* gene subsets, and can be adapted to predict sequence and domain compositions from tags of other variable gene families.

## Acknowledgements

We thank MalariaGen. TO is supported by the Wellcome Trust grant 104111/Z/14/ZR.

