## Supplemental material for "Varia: Prediction, analysis and visualisation of variable genes"

**Supplementary data of**
**Varia: Prediction, analysis and visualisation of variable genes**

Gavin Mackenzie[1], Rasmus W. Jensen[2], Thomas Lavstsen[2,*] and Thomas D. Otto[1,*]
[1]Centre of Immunobiology, Institute of Infection, Immunity & Inflammation, MVLS, University of Glasgow, Glasgow, UK. [2] Centre for Medical Parasitology, Department of Immunology & Microbiology, University of Copenhagen and Department of Infectious Diseases, Rigshospitalet, 1017 Copenhagen, Denmark.

**Content**

1. Basic information on *var* gene and PfEMP1 domain composition useful for understanding and interpretation of Varia design and output

2. *Var* gene and annotation databases

3. Varia pipelines
   3.1 *Var* Identification and Prediction (Varia_VIP) pipeline
   3.2 Gene Expression Module (Varia_GEM) pipeline

4. Varia parameter testing

5. Examples of Varia use

### 1. Basic information on *var* genes and PfEMP1 domain composition useful for understanding and interpretation of Varia design and output

*Plasmodium falciparum* depends on sequestration of infected erythrocytes in host tissue. Several different human receptor binding phenotypes have evolved, mediated by the parasite-derived *P. falciparum* membrane proteins 1 (PfEMP1) adhesion molecules exported to the infected erythrocyte surface(Jensen, et al., 2020). PfEMP1 are targets of acquired immunity, and in response, the protein family has expanded and diversified to each parasite genome containing ~60 copies of PfEMP1-encoding *var* genes of 6-12kb in length(Otto, et al., 2018). These genes differ in sequence within and between parasites but collectively bestow each parasite a similar repertoire of PfEMP1 molecules and phenotypes. The extracellular part of PfEMP1 (encoded by the *var* exon1) is composed of multiple DBL (Duffy binding like) and CIDR domains capable of binding specific human receptors. DBL and CIDR domains are classified into few ***main*** domain types, DBLα- ζ and CIDRα-γ, which have been further divided into domain ***subtypes*** according amino acid similarity (Rask, et al., 2010). Due to varying sequence diversity homogeneity between main domain types, the stringency by which the domain subtypes are defined, differ between main domain types. A comprehensive description of PfEMP1 domain subtyping can be found in (Otto, et al., 2019). The domain composition of each PfEMP1 can vary but follows a general pattern (Figure S1) which includes an N-terminal NTS-DBLα-CIDR “head structure” followed by varying combination of domains in a semi-conserved order of main domain types. The exception to this rule is the highly conserved *var2csa* and *var3* genes, which encode atypical domain compositions. The anchoring intracellular part of PfEMP1 is encoded by the *var* exon2.

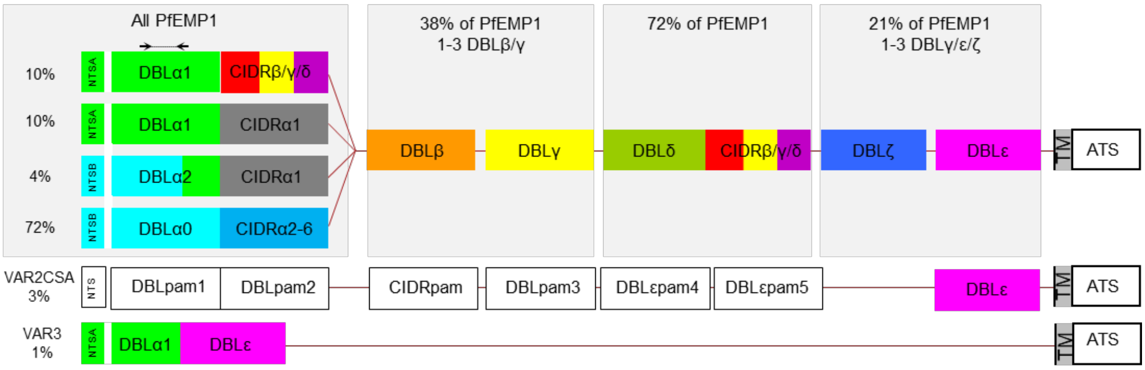

**Figure S1:** Overview of *var* gene-encoded PfEMP1 domain structures and the common distribution of different types and domain compositions of *var* genes in the average *P. falciparum* genome. Arrows indicate location of commonly used annealing sites for degenerate primers, which can target the genes to generate DBLα sequence tags.

PfEMP1 phenotypes can to some extend be predicted from sequence. The VAR2CSA PfEMP1 binds placental chondroitin sulphate A(Salanti, et al., 2003), and placental sequestration of parasite-infected erythrocytes expressing VAR2CSA is the cause of placental malaria(Salanti, et al., 2004). The N-terminal CIDR domain type predicts mutually exclusive binding phenotypes of all other PfEMP1. Thus, CIDRα1 domains bind EPCR, with the exception of CIDRα1.2 and CIDRα1.3 subtypes found in the *var1* pseudogenes(Turner, et al., 2013), and all CIDRα2-6 domain variants are considered CD36-binders(Hsieh, et al., 2016). ICAM1 binding is linked to subsets of DBLβ domains, often, but not always, of the DBLβ3 and DBLβ5 subtypes(Janes, et al., 2011; Lennartz, et al., 2017). The function of other DBL and CIDR variants is unclear. Although other PfEMP1 functions including binding to PECAM1, HABP1 IgM, A2M have been proposed, these have not been fully defined molecularly(Hviid and Jensen, 2015).
A large body of evidence has linked parasites expressing EPCR-binding PfEMP1 to development of severe malaria symptoms(Bernabeu and Smith, 2017), but little is known about the relative contribution of other PfEMP1-mediated host interactions to clinical outcomes of infection. Analysis of *var* gene expression in patient samples is difficult due to the diversity of the genes. Although it is possible to obtain full-length *var* genes by assembling data from full genome sequencing, assembly and expression profiling from RNA sequencing is error prone, expensive and often challenging due to the small volumes of blood, which can be drawn from severely ill children(Andrade, et al., 2020). A cost-effective alternative has been to sequence RT-PCR-amplified DBLα tags, and use these to quantify the relative expression level of *var* genes in a patient sample. However, until now it has not been possible to infer the domain composition of the entire encoded PfEMP1 from the short sequence tags.

DBL domains are composed of alternating variable and conserved blocks. Universal primers targeting homology blocks in DBLα domains can be used to relatively unbiased amplify *var* expression tags from cDNA to infer the relative distribution of different *var* genes expressed in a patient (Bull, et al., 2005). The function of the DBLα domain itself is unknown, but as the DNA sequence diversity in the DBLα-tag region is extensive, it is possible that the flanking sequence of the originating genes can be predicted from the DBLα-tag, if sufficient information on global sequence diversity is available. One important caveat however, is that *var* genes exhibit recombination hotspots within DBLα domains and mid-*var* around the 5´end of DBLδ or DBLγ domains(Rask, et al., 2010). The historic recombination events may impede prediction of full-length *var* gene sequences. Moreover, prediction of the exact sequence of the intracellular membrane-anchoring domain encoded by *var* exon2 is unlikely to be possible as the highly similar intron sequences compromise correct assembly of exon1 and exon2(Otto, et al., 2019). With these features in mind, we developed the Varia tool for prediction of the *var* gene sequence flanking short sequence tags of *var* genes with particular focus on DBLα tags.

**2. *Var* gene and annotation databases**

Varia uses a *var* gene database to search for sequence tag hits. The Varia database is composed of two datasets from clinical isolates: (1) 2400 sequences from the Pf3K community resource (https://www.malariagen.net/projects/Pf3k) (2) the *P. falciparum* Community Project (version January 2016, https://www.malariagen.net/projects/p-falciparum-community-project), which comprises the Pf3k community resource. The *var* genes from the Pf3K dataset were assembled in (Otto, et al., 2019), and were taken from ftp://ftp.sanger.ac.uk/pub/project/pathogens/Plasmodium/falciparum/PF3K/varDB/FullDataset/ or https://github.com/ThomasDOtto/varDB/ Datasets/FullDataset. The further 755 isolates from the Community Project were assembled and annotated for this project, using the *var* gene assembly pipeline as described in (Otto, et al., 2019) and the sequences were added to https://github.com/ThomasDOtto/varDB/tree/master/Datasets/Additional755/.

As we are interested in the prediction of sequences and domains, sequences shorter than 3kb were excluded, resulting in 162,827 *var* genes from the Pf3K dataset and 41,375 from the additional sequences from the Community project. This dataset was complemented with 1,393 full length *var* genes from 15 PacBio reference genomes(Otto, et al., 2018), including the canonical reference 3D7(Bohme, et al., 2019). These sequences were obtained from long read assemblies and manually curated and therefore used as “truth set” for testing. In total the complete database contains 205,595 *var* gene sequences.

It should be noted that not all of the genes from clinical isolates are full-length *var* genes and in general, the exon2 is missing from these genes. From translated amino acid sequences, we predicted domain subtypes described in (Rask, et al., 2010) using HMMer models developed in (Otto, et al., 2019). We used the command “hmmscan --cpu 12 --noali -E 1e-6 --domE 1e-6” and parsed the results with in-house Perl script. The Perl script and the HMMer models for the prediction are available at https://github.com/ThomasDOtto/varDB/.

As annotation was performed on complete and incomplete *var* sequences, these criteria may leave some sequence ends with incomplete domains unannotated.

**3. Varia pipelines**

Varia is based on Linux, programmed in Bash and Python and is freely available on https://github.com/GCJMackenzie/Varia/. Varia builds on several basic bioinformatics tools, like NCBI blast, mcl, *Circos* and Samtools. Please see the GitHub page for a complete list of pre-requisites. It was tested in Linux and Mac environments.

Varia has two pipelines (Figure S2), the *var* identification and prediction (Varia_VIP) and the Gene Expression Module (Varia_GEM).

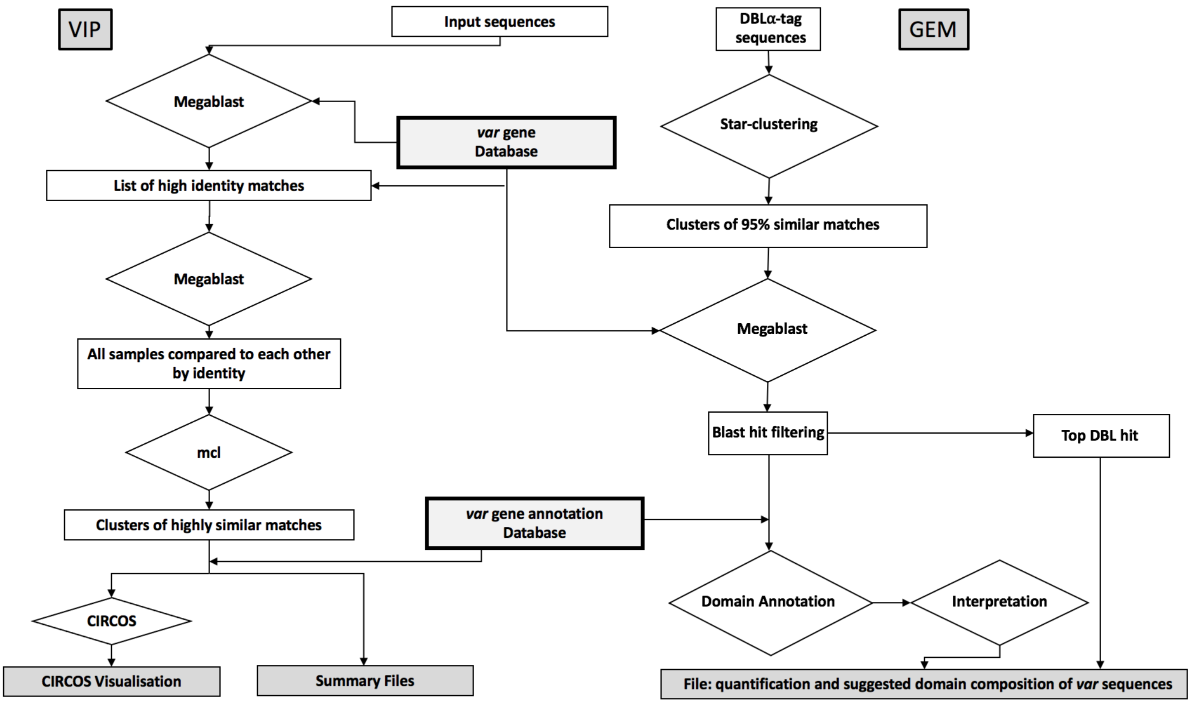

**Figure S2. Flowchart of Varia pipelines.** Filled rectangles indicate output files; diamonds indicate tools and data processing.

#### 3.1 *Var* Identification and Prediction (Varia_VIP) pipeline

*Arguments*
Varia_VIP is run using the following command line:

Varia.sh VIP [optional arguments] -i [input tag file]

-i is the only mandatory argument required to run Varia_VIP as this specifies the input file to be used. Varia_VIP also has a number of optional arguments, which can be used to change the output directory and change various filters used throughout the module, a detailed list of these options and their default settings can be found in the readme file, or by using:

Varia.sh VIP -h

*Database choice*
The database used by Varia_VIP is selected as part of the installation script. As default the program is set to use the provided paired "*var* sequence" and "*var* annotation" databases. Once these databases are downloaded, execution of the installation script, Install_Varia.sh, will prepare the database files for downstream analysis (generating the search indexes). The default database should be downloaded from https://github.com/ThomasDOtto/varDB/tree/master/Datasets/Varia/.

The *var* sequence and *var* annotation database files will be soft-linked into the Varia domains subdirectory. This enables the user to use other databases (e.g. to apply Varia to other variable gene families). To do this, the sequence database should be in fasta file format, and an accompanying domain annotation file containing matching entries should be formatted using formatdb from the NCBI blastall package. The user can change databases used by Varia by deleting soft links to the fasta index and database files, then rerunning the installation script.

*Environment setup and input data*
Input sequence tags should be provided to Varia_VIP in fasta file format. A batch of multiple fasta-formatted sequence tags can be submitted in a single input file and Varia_VIP will run each tag individually. Each tag must be longer than 200 bps (default setting) as this minimum length is later used to filter the database blast results.

As input parameters, Varia_VIP requires the name of the input tag file. In addition, the user can modify several parameters, such as the *identity cut-off* used for filtering database hits. After checking the validity of parameters given, a new result directory [name of input file]-[identity filter]-Varia_Out is generated (name and location can be changed with the -o parameter).

*Search for tag-related sequences in the var sequence database*
After the environment is set up, the input file is duplicated into a temporary file to allow for modifications without altering the original file. The first input tag is extracted from the input file, placed in a temporary file and compared to the sequence database using Megablast (Morgulis, et al., 2008). If there are no database hits, VARIA_VIP will move onto the next input tag. For each tag with a database hit, the output blast file is filtered to collect the names of all database hits that are longer than the "database length filter" (-l parameter, default 200 bps) and higher than the "database identity filter" (-f parameter, default 99%).

*Comparison and clustering of hit sequences*
Next, the names of the filtered hits and the *samtools faidx* tool are used to create a new fasta file containing the sequence for each hit. *Formatdb* is used to temporarily make this fasta file into a database that Megablast can use in a blast-all-against-all search to generate an output file containing every region of alignment between each gene and every other gene in the fasta file. These results are filtered to include all results with sequence identities higher than the "self-blast identity filter" (-c parameter, default 99%) over the percentage of length set by the "self-blast length filter" (-p parameter, default 80% of either the aligned sequences length). The sequence name and alignment length are extracted from each filtered hit and formatted for Markov Clustering (*mcl*) of the sequences into clusters of near identical sequences. Due to high sequence variation between *var* genes, it is recommended to use the default self-blast settings of 99% identity and >80% length of either sequence criteria to ensure members of a cluster are highly similar both in sequence and domain composition. The largest sequence in each cluster is then used as a representative of the cluster, for example in the circus plots. Clusters are numbered according to number of hit sequences included, i.e. cluster 1 contains most hits.

*Plot generation*
The representative sequence of all clusters relating to a specific query tag, and their domain composition extracted from the annotation database, are then formatted for visualization in *Circos* plots. *Circos* allows multiple plots to be shown for each cluster. Annotation pertaining to each sequence is presented around the circumference of the visualisation. Figure S3 shows different types of information annotated to an individual sequence in different tracks (A-E) radiating out from the centre of the entire visualisation. Varia provides several input files for track labelling and other features. The display of the different features is stored in the "Varia.conf" *configuration file,* see below.

All input files needed to generate the plots are retained by Varia to allow users to customise the plot of a single sample.

Configuration File: Varia_VIP uses a standard configuration file for all *Circos* plots. This file details what tracks are present, the input files to be used for them, the positioning of tracks and labels and formatting of text size and font etc. Given the large number of parameters, it is recommended that users edit a copy of the provided Varia configuration file if should they wish to alter the plot. Directly editing the configuration file will alter all plots Varia makes from then on. The configuration file “Varia.conf” is found in the “Varia1_5/scripts/” directory.

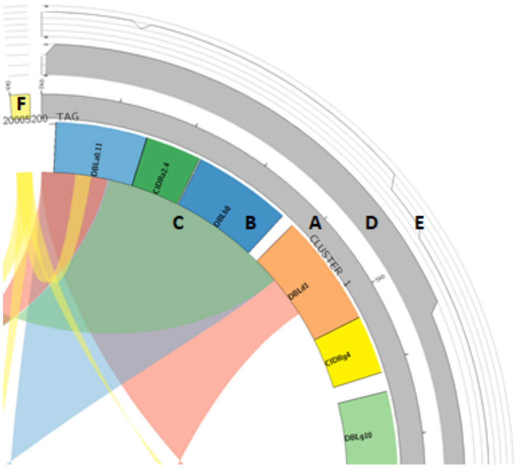

**Figure S3: Explanation of *Circos* output on the example of** cluster 1 of PfIT_020005200 (DBL**α** domain): A: Chromosome track, displaying cluster name and length of largest gene in the cluster. B: Domain plot showing labelled domain composition colour coded. C: Ribbon plot of regions of identity shared by the clusters, coloured to help distinguish ribbons. D: Coverage plot quantifying how many genes within the cluster share that region with the largest sequence. E: Coverage plot quantifying the number of clusters that share that region of the genes sequence to other genes in the comparison (similar to C).

Chromosome Track: Grey track labelled A on Figure S3. The chromosome track is the only mandatory track in a *Circos* plot. Each cluster-representative sequence is labelled by its cluster number and acts as an x-axis for other annotation plots relating to the cluster. Ticks represent 1000 bp intervals. **F** on Figure S3 represents the input tag used and is high-lighted in yellow.

Domain track: Labelled B on Figure S3. Track B shows the encoded PfEMP1 domain composition of the cluster-representative sequence. Each block on the plot represents a domain and shows its size and positioning along the length of the largest sequence. These blocks are labelled based on the sequence annotation database selected during installation, the soft link to the file *vardb_domains.txt,* which can be found in the domains directory.

The colour of the blocks is determined based on the domain name and what colour is assigned to it in the colour map file *domain_color_map.txt* in the “domains” directory. The user can change the colours by editing this file.

Ribbon track: Labelled C on Figure S3. The ribbon plot is used to show regions that share at least 200bp and an identity ≥ 99% between the largest sequences of other clusters. Each ribbon will show the position and length of regions that are nearly identical between two cluster-representative sequences. The colour of links cycles between red, blue and green to help differentiate the ribbons on a plot and show overlapping ribbons matching to the same place. Ribbons between the input tag (F) and other clusters, are shown in yellow. There should always be a link between the input tag and every other cluster.

Inter-cluster coverage track: Labelled E on Figure S3. This plot is a quantification of the information shown on the ribbon plot. Here, the Y axis shows the number of plotted sequences that share identical across a region of 100 base pairs. To obtain this, an array of integers is set up for each base pair position in the sequence.

For each position, the ribbon plot file is searched for matches to the plotted sequence, and when a match is found, 1 is added to each base pair position within the range of the ribbon. Once the ribbon plot file has been fully searched, the array is split into 100bp segments and the median is calculated for that segment. The largest coverage value for the entire plot is then set as the maximum Y value for the entire plot. Finally, files are generated to display the Y axis and the maximum and minimum values.

Intra-cluster coverage track: Labelled D on Figure S3. Similar to the inter-cluster coverage plot, this plot shows the regions of matching sequence shared between genes contained within the cluster. The genes in each cluster are compared to the largest gene in that cluster and the results are mapped into an array as detailed above and the median is found for each 100 bp segment. Max value is found, and files are generated to show the Y axis.

*Summary Files*

For each input sequence tag, summaries of the extracted database hit sequences are also returned in excel-readable tables.

Cluster summary (Figure S4)**:** The *cluster summary* file shows information for every gene that was clustered by mcl, as a separate entry. Each entry shows the cluster number and name, the name of the gene, the length of the gene in base pairs, the country where the gene was isolated and the encoded domain composition. If specific information points are lacking, "n/a" is reported. An example can be found on the GitHub site, in the example directory.

Final summary (Figure S5)**:** The *final summary* file shows information on each cluster-representative sequence returned by Varia_VIP. Each entry in the file shows: cluster name, the number of genes in the cluster, the name, sequence length and encoded domain composition of the longest gene (the cluster-representative sequence) in the cluster and, the number of different countries reported in the cluster. An example can be found on the git-hub, under Example/VIP/output.

| Cluster_ID | Seq_ID | Seq_length | Country_Distribution | Subdomains |
|---|---|---|---|---|
| 1 | PH0444-C.g34 | 5035 | Cambodia | NTSA3 DBLa1.2 CIDRa1.5 DBLg17 DBLd5 |
| 1 | PH0194-C.g145 | 8319 | Cambodia | NTSA3 DBLa1.2 CIDRa1.5 DBLg17 DBLd5 CIDRb3 DBLb7 DBLg9 |
| 1 | PH0241-C.g416 | 8262 | Cambodia | NTSA3 DBLa1.2 CIDRa1.5 DBLg17 DBLd5 CIDRb3 DBLb7 DBLg9 |
| 1 | PH0820-C.g184 | 8262 | Cambodia | NTSA3 DBLa1.2 CIDRa1.5 DBLg17 DBLd5 CIDRb3 DBLb7 DBLg9 |
| 1 | PH0823-C.g23 | 8319 | Cambodia | NTSA3 DBLa1.2 CIDRa1.5 DBLg17 DBLd5 CIDRb3 DBLb7 DBLg9 |
| 1 | PH0824-C.g47 | 8262 | Cambodia | NTSA3 DBLa1.2 CIDRa1.5 DBLg17 DBLd5 CIDRb3 DBLb7 DBLg9 |
| 1 | PH0839-C.g640 | 8262 | Cambodia | NTSA3 DBLa1.2 CIDRa1.5 DBLg17 DBLd5 CIDRb3 DBLb7 DBLg9 |
| 1 | PC0016-C.g67 | 8262 | Kenya | NTSA3 DBLa1.2 CIDRa1.5 DBLg17 DBLd5 CIDRb3 DBLb7 DBLg9 |
| 1 | PH0152-CW.g96 | 8214 | Cambodia | NTSA3 DBLa1.2 CIDRa1.5 DBLg17 DBLd5 DBLb7 DBLg9 |
| 1 | PH0390-C.g5129 | 8262 | Cambodia | NTSA3 DBLa1.2 CIDRa1.5 DBLg17 DBLd5 CIDRb3 DBLb7 DBLg9 |
| 1 | PH0819-C.g583 | 3147 | Cambodia | NTSA3 DBLa1.2 CIDRa1.5 DBLg17 |
| 1 | PD0863-C.g901 | 7077 | Thailand | NTSA3 DBLa1.2 CIDRa1.5 DBLg17 DBLd5 CIDRb4 DBLb9 |
| 1 | PH0291-C.g8 | 8147 | Cambodia | CIDRa1.5 DBLg17 DBLd1 CIDRb3 DBLb7 DBLg9 |
| 2 | PV0303-C.g485 | 6023 | Vietnam | NTSA3 DBLa1.2 CIDRa1.5 DBLb3 DBLd1 CIDRb5 |
| 2 | PH0332-C.g290 | 6444 | Cambodia | NTSA3 DBLa1.2 CIDRa1.5 DBLb3 DBLd1 CIDRb5 |
| 2 | PH0545-C.g745 | 6444 | Cambodia | NTSA3 DBLa1.2 CIDRa1.5 DBLb3 DBLd1 CIDRb5 |
| 2 | PH0555-C.g426 | 6444 | Cambodia | NTSA3 DBLa1.2 CIDRa1.5 DBLb3 DBLd1 CIDRb5 |
| 2 | PH0556-C.g464 | 6444 | Cambodia | NTSA3 DBLa1.2 CIDRa1.5 DBLb3 DBLd1 CIDRb5 |
| 2 | PH0557-C.g457 | 6444 | Cambodia | NTSA3 DBLa1.2 CIDRa1.5 DBLb3 DBLd1 CIDRb5 |
| 2 | PH0574-C.g356 | 6444 | Cambodia | NTSA3 DBLa1.2 CIDRa1.5 DBLb3 DBLd1 CIDRb5 |
| 2 | PH0717-C.g208 | 6444 | Cambodia | NTSA3 DBLa1.2 CIDRa1.5 DBLb3 DBLd1 CIDRb5 |
| 2 | PD0476-C.g308 | 6501 | Thailand | NTSA3 DBLa1.2 CIDRa1.5 DBLb3 DBLd1 CIDRb5 |
| 2 | PD0492-C.g677 | 6444 | Thailand | NTSA3 DBLa1.2 CIDRa1.5 DBLb3 DBLd1 CIDRb5 |

**Figure S4** Example of Varia_VIP output showing the beginning of the cluster summary file for input tag PfDd2_010005500:532-814. Columns from left to right: The cluster number each entry is assigned to; the sequence ID of the Varia database hit sequence; the length in base pairs of the hit sequence; the sequence country of origin; and the domain subtype composition of the hit gene.

| Cluster_name | Cluster_size | Longest_seq | length | 80%_matches | Coutry_distrib | Subdomains |
|---|---|---|---|---|---|---|
| Cluster 1 | 13 | PH0194-C.g145 | 8319 | 7 | 3 | NTSA3 DBLa1.2 CIDRa1.5 DBLg17 DBLd5 CIDRb3 DBLb7 DBLg9 |
| Cluster 2 | 11 | PD0476-C.g308 | 6501 | 10 | 3 | NTSA3 DBLa1.2 CIDRa1.5 DBLb3 DBLd1 CIDRb5 |
| Cluster 3 | 7 | PD0872-C.g203 | 6813 | 6 | 1 | NTSA3 DBLa1.2 CIDRa1.5 DBLg11 DBLd1 CIDRb5 |
| Cluster 4 | 3 | PV0113-C.g13 | 10608 | 0 | 2 | NTSA3 DBLa1.2 CIDRa1.5 DBLg17 DBLd5 CIDRb3 DBLb7 DBLg9 DBLd1 CIDRb1 |
| Cluster 5 | 2 | PH0151-CW.g22 | 8229 | 1 | 1 | NTSA3 DBLa1.2 CIDRa1.5 DBLg7 DBLd5 CIDRb3 DBLb9 DBLg9 |
| Cluster 6 | 2 | PD0512-C.g297 | 7665 | 0 | 1 | NTSA3 DBLa1.2 CIDRa1.5 DBLb3 DBLg11 DBLd1 CIDRb6 |
| Cluster 7 | 2 | QG0236-C.g2368 | 6885 | 1 | 1 | NTSA3 DBLa1.2 CIDRa1.5 DBLb3 DBLd1 CIDRb1 |
| Cluster 8 | 1 | PT0257-Cx.g2301 | 8274 | 0 | 1 | NTSA3 DBLa1.2 CIDRa1.5 DBLg17 DBLd5 CIDRb3 DBLb6 DBLg11 |
| Cluster 9 | 1 | PF0819-C.g2715 | 7742 | 0 | 1 | NTSA3 DBLa1.2 CIDRa1.5 DBLg17 DBLd5 CIDRb3 DBLb6 |
| Cluster 10 | 1 | PH0141-CW.g1 | 8961 | 0 | 1 | DBLa0.3 CIDRa1.5 DBLg11 DBLd5 CIDRb4 DBLe11 DBLz2 |

**Figure S5** Example of Varia_VIP output showing the final summary file for input tag PfDd2_010005500:532-814. Columns from left to right: The cluster number; the number of sequences in the cluster; the sequence ID of the longest database hit sequence in the cluster i.e. the cluster-representative sequence; the length in base pairs of the longest hit sequence; number of sequences in the cluster that are 99% identical over at least 80% of the length to the largest sequence in the cluster; the number of different countries of origin in the cluster; the domain subtype composition of the cluster-representative sequence.

### 3.2 Gene Expression Module (Varia_GEM) pipeline

*Environment setup and input data*

One or more input FASTA files containing DBLα tag sequences are required for Varia_GEM to run. The parameter are similar to the VIP module with the addition of:

> -c - *min_cluster_size* is specified in the command line, the program will exclude downstream analyses of sequences representing clusters containing less than the *min_cluster_size* number of sequences. Default *min_cluster_size* setting is 10 sequences per cluster.

The program creates temporary fasta files, where each sequence is written and for each fasta file, Varia_GEM creates a subfolder for the output files generated downstream.

*Clustering of sequences and search for tag-related sequences in the var sequence database*
For each fasta file, DBLα tag sequences are clustered into groups sharing at least 95% nucleotide identity across minimum 200 bps using *Vsearch* and written to separate cluster files in the fasta file subfolder, numbered from 1 to n. One sequence from each cluster is then used to search for genes with similar DBLα tag sequences in the *var* gene database using Megablast. Megablast hits are filtered keeping E-value <1e-2 and capped at maximum 200 hits. The DBLα domain subtype most prevalent among the top 50 hits is determined, and all blast hits are filtered keeping only hits with >95% identity across minimum 200 nucleotides to the DBLα-tag. The name of each filtered hit gene is used to retrieve the encoded domain composition of the hit gene from the domain annotation database ("vardb_GEM_domains.txt").

*Predicting the domain composition of the gene from which the DBlα-tag was derived*
For each hit gene, the domain type found at each domain position 1-10 (D1-D10) (the DBLα domain is at domain position 2, see Figure S1) is logged and the summarized counts for all hit genes are written to a fasta file-specific excel sheet named with the input fasta file name. The counts of different domain types found at same domain position is processed in an interpretation step, in which the main domain type at each position is called if a type accounts for more than 66% of the total domain count at the position. If no consensus is found, no domain annotation is made. If a consensus is found, the program checks if there is consensus on the domains subtype classification by checking if a domain subtype within the main domain type dominates with more than 66% of the counts. If no domain subtype consensus is reached, only the main domain type is given. For example, if 7 of 10 hit genes encode a DBLβ domain at position D4, and five of these are of the DBLβ12 subtype, the domain is called DBLβ12. If 7 of 10 hit genes encode a DBLβ domain at position D4, and four of these are of the DBLβ12 type, the domain is called DBLβ. All *var* genes (except *var3* not amplified by commonly used DBLα tag primers (Lavstsen, et al., 2012)) contain domains at position D1-D5, but only some have domains at position D6-D10. By including the criteria that annotation of domains at D6-D10 was only called if based on >40% the hit genes, we found that over-prediction of non-existing domains could be reduced to occur in <10% (data not shown).

*Excel result sheet output*
Each run of Varia_GEM will result in one excel file containing one sheet per input fasta file as well as one sheet termed "All_samples_summary" containing a summary of all of fata file analyses.
In the fasta-file-specific sheets, the predicted consensus domain compositions are written as binary counts in an interpretation section of the sheet and as domain positional and non-positional data along with a calculation of the proportion of genes or transcripts within the input fasta file that encode a given domain type. The consensus domain composition is also written in the fasta file-specific excel sheet as a domain string for each tag. The sheet also contains the *Sample_ID*, *Cluster_size*, the cluster-representative sequence, the most frequent DBLα type among top fifty blast database hits and the *#hits in varDB*.
The "All_samples summary" sheet contains the summarized relative expression level for each main domain type found in position D1-D10 and all main domain types and subtypes for all fasta files analysed. The data is given in a table format allowing subsequent merging with phenotypic or clinical data pertaining to each fasta input file (i.e. patient sample).

An example of the output of both modules can be found in the example directory on the GitHub repository.

**4. Varia parameter testing**
Varia_VIP can analyse sequence fragments from any part of the *var* exon 1. In practice *var* gene tags are likely to be generated from sequencing of PCR-amplified fragments spanning across variable regions roughly corresponding to the subdomain 2 of DBL domains. The varying sequence diversity homogeneity between main

types of DBL domains is likely to affect the ability of Varia_VIP to predict correct domain compositions. To assess this, Varia_VIP predictions were made from DBL tags derived from 971 DBLα-ζ domains (Table S1). These sequences are from the curated reference dataset and therefore seen as a truth set. For the running of the pipeline, these sequences were excluded from the database. The cognate genes were extracted from the database and the tags were run using an identity filter threshold of 99%, 95% and 90% over 150 bps (DBL Subdomain 2 domain lengths vary between DBL types with DBLε domains being shortest).

**Table S1. Analysis of database hits and gene predictions from Varia_VIP analysis of DBLα-ζ tags.** Sample DBL tags 971 different DBLα-ζ domains were run through Varia_VIP using a length filter of 150 base pairs and an identity filter of 99%, 95% and 90%. The hit rate shows the proportion of queried tags that had one or more hits in the *var* gene database. The average number of clusters into which hit genes were grouped and the proportion of genes for which a correct domain subtype annotation was found in any cluster, or in the five clusters with most hit sequences (top 5), is shown. Also shown is the proportion of genes for which a sequence matching the reference gene by 99% identity over at least 80% of the full sequence, was found among the top five Varia_VIP clusters.

| Domain | No. tags tested | Hit rate (%) | | | Average No. of clusters | | | Percentage correctly annotated genes (any cluster) | | | Percentage correctly annotated genes (top 5 Clusters) | | | Percentage perfect DNA sequence hits (top 5 clusters) | | |
|---|---|---|---|---|---|---|---|---|---|---|---|---|---|---|---|---|
| Seq. ID threshold→ | | 99% | 95% | 90% | 99% | 95% | 90% | 99% | 95% | 90% | 99% | 95% | 90% | 99% | 95% | 90% |
| DBLα | 293 | 95 | 99 | 100 | 14 | 34 | 92 | 72 | 75 | 78 | 66 | 62 | 53 | 29 | 27 | 22 |
| DBLβ | 127 | 94 | 96 | 97 | 13 | 41 | 100 | 73 | 73 | 73 | 69 | 65 | 53 | 39 | 35 | 27 |
| DBLδ | 256 | 96 | 99 | 100 | 15 | 34 | 60 | 74 | 74 | 77 | 69 | 63 | 55 | 33 | 29 | 26 |
| DBLγ | 138 | 91 | 93 | 93 | 18 | 61 | 146 | 71 | 70 | 71 | 64 | 55 | 44 | 41 | 36 | 24 |
| DBLε | 109 | 78 | 78 | 78 | 21 | 61 | 120 | 54 | 56 | 56 | 51 | 44 | 39 | 25 | 17 | 13 |
| DBLζ | 48 | 96 | 100 | 100 | 29 | 99 | 263 | 60 | 63 | 63 | 54 | 38 | 29 | 35 | 21 | 10 |

This showed that tags from DBLα, β, δ and γ domains resulted in fewer clusters and more frequent correct annotations compared to DBLε and DBLζ domains. This observation can be explained by the differences in the distribution of diversity within these main domain types. DBLα, β, δ and γ domains are highly diverse in sequence and diversity is homogeneous. Conversely, DBLε and DBLζ domain sequences distribute into distinctly different groups of highly similar sequences. Thus, as identical DBLε and DBLζ domains are found in many different PfEMP1 variants, correct predictions are difficult to make from these domain tags.

Next, we focussed on the DBLα-tag. This is the most commonly analysed *var* sequence tag as it is found in all *var* genes but is highly diversified and can be targeted by universal primers (not targeting the highly diverse *var2csa* and *var3* genes) (Bull, et al., 2005; Lavstsen, et al., 2012). One or more hits were found for 95%, 99% and 100% of the tested DBLα tags, when searching with identity filter of 99%, 95% and 90%, respectively.
A cluster-representative sequence with domain subtype annotation perfectly matching the query gene was found for 72%, 75% and 78% of queries, when filtering hit sequences by 99%, 95% or 90% identity, respectively. We also assessed correct gene predictions defined as DNA sequences with 99% shared identity to the reference gene sequence for 80% of the length. Correctly predicted gene sequences were found for 29%, 27% and 22% of the query genes, when database hits were filtered for 99, 95 or 90 % identity to the DBLα tag. This revealed that correct predictions were most often found among the Varia_VIP clusters containing most database hits. Allowing more diversity in the DBLα tags, and thereby higher number of hit sequences increased likelihood of finding one correct domain prediction. However, this came at the cost of the correct sequence and domain annotation not placed in the clusters with most sequence hits.

For high-throughput expression analysis of *var* DBLα tags in Varia_GEM, a single likely domain composition for each query tag must be predicted. To further assess and determine parameters for optimal coverage and accuracy of domain predictions by Varia_GEM, we tested main domain type and domain subtype predictions at different identity threshold values. We used 220 DBLα0, 1 and 2 tag sequences (660 in total), randomly selected from the *var* gene database. The genes were removed from the database, and the database was searched for genes sharing 99, 97, 95, 93% and 85% identity across 200 bp of the DBLα-tags. This resulted in an average ~25 hits per DBLα tag at 99% threshold increasing to 44 at 93% and to 114 at 85% (Figure S6A). The consensus domain composition generated for each DBLα-tag was compared to known domain composition of its cognate gene to calculate the accuracy of the predictions at each position of the domain (Figure S6B). The accuracy was calculated both by the main domain type and domain subtype level, as follows:

a) Average main domain accuracy: An annotation was considered incorrect if the predicted domain main type was wrong or if a *non-existing* domain was predicted (i.e. a domain was predicted at a position where no domains were present in the original sequence).

b) Average domain subtype accuracy: An annotation was considered incorrect if the predicted domain subtype was *wrong* or if a *non-existing* domain was predicted (i.e. a domain was predicted at a position where no domains were present in the original sequence).

Next, the frequency by which the tool could predict the correct *var* gene domain compositions (coverage) was calculated as:

c) Average coverage (main domain type): The proportion genes with a correctly predicted main domain type at said position. A domain annotation was considered incorrect if a *wrong* main domain type was predicted or a domain annotation was missing (i.e. no domain type was predicted where a domain was present in the original sequence).

Lowering the blast identity threshold and thereby including a higher number of hits, resulted in fewer predictions and fewer correct domain annotations. However, main domain predictions were robust, with 85% accuracy over first 1-4 domains. Lowering the blast identity threshold from 99% to 93% caused a slight decrease in main type accuracy, but a moderate decrease in coverage and domain subtype accuracy. Lowering the identity threshold further to 85% caused a significant drop in main domain coverage and domain subtype accuracy, but again only a small decrease in main domain type accuracy. The most significant effects from lowering the blast identity threshold, was seen on domains at position D5-8. The level of coverage and accuracy was similar for DBLα0, 1 and 2 tag sequences (not shown).
These observations reflect the nature and the classification of *var* genes well. Main domain types (DBLα-ζ, and CIDRα-γ) are well defined, but the sequence diversity within the main domain types differs between main domain types and domain subtyping is uncertain (Otto, et al., 2019; Rask, et al., 2010). For example, found at domain position 2, CIDRα domain sequences separate distinctly into two groups, CIDRα1 or CIDRα2-6 domains, but the 18 subsets of CIDRα2-6 domains, all expected to bind CD36, are less well segregated and defined. Also, the recombinogenic nature of *var* genes make it likely that gene and domain predictions will be most accurate nearest to the analysed tag. This explains the drop in main domain coverage and domain subtype accuracy with the domain position, and the significant drop around domain position D5, which corresponds to the main mid-*var* gene recombination hotspot (Figure S6B). Encouragingly, the overall main domain accuracy across all domain positions, shows that if a main domain type is predicted, this is very likely to be true.

Most frequent and correct domain predictions were generated using the 99% blast identity threshold. However, this threshold is likely to be too stringent for analysis of sequencing data, which may contain various minor sequence errors. Instead, a 95% threshold was chosen as default for Varia_GEM. This threshold results in on average a 0-10% drop in main domain type coverage and domain subtype accuracy, and a 0-3% decrease in main domain type accuracy. Using the 95% identity threshold, the effect of the number of blast hits on domain prediction was investigated (Figure S6C). This showed that a higher number of hits adversely affected the proportion of correct predictions (coverage) but did not affect the accuracy of the predictions, and that including predictions made based on only 1 hit sequence results in most frequent correct predictions. Based on this, Varia_GEM was set to predict on every tag regardless of number of database hits.

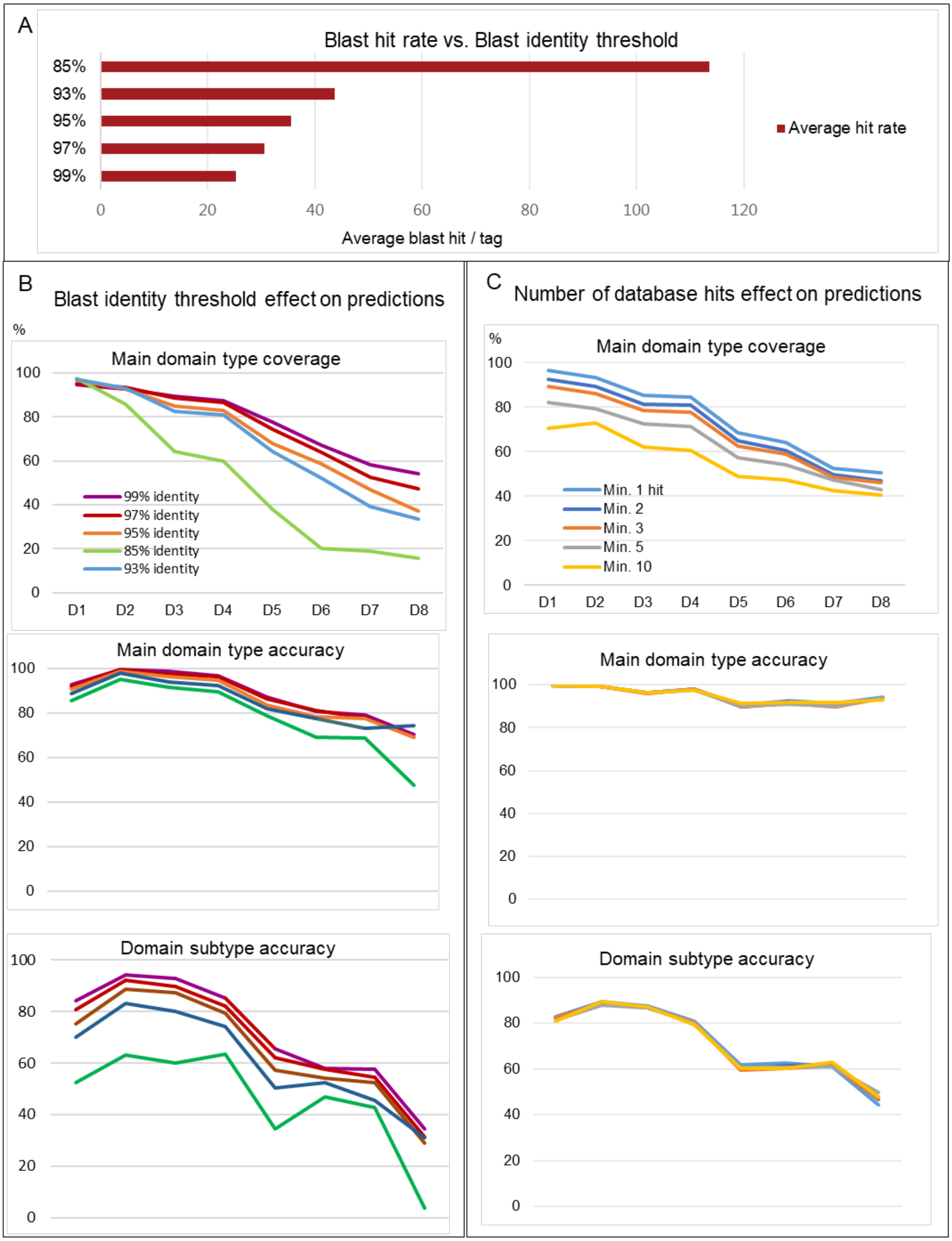

**Figure S6. Analysis of database hits and gene predictions by Varia_GEM analysis of DBLα tags.** A) Average blast hit per query DBLα tag sequence at different blast identity thresholds. B) Plots showing main domain type coverage and domain type accuracy of predictions across the first eight domains (D1-D8) for 660 DBLα tags extracted from genes encoding 220 DBLα0, 1 and 2 domains. Data shown for different blast identity thresholds (99-85%). C) Domain prediction coverage and accuracy of predictions made at 95% blast identity threshold shown by number of database hits.

Users should note that the current *var* gene database is biased towards genomes from South East Asian parasite isolates (Otto, et al., 2018). When testing tags from 15 reference isolates of different origin, 94% of tags from African isolates had one or more database hit with over 95% sequence identity over 200 base pairs, compared to 99% of DBLα tags from Asian isolates. This resulted in a correctly annotated cluster-representative sequence by Varia_VIP (any cluster) for 56% vs. 76% of the query tags from African vs. Asian isolates, respectively.

### 5. Examples of Varia use

*Example input and output of Varia_VIP*

Useful output for the user in the Varia_VIP are the tables (Figure S4 and S5) and the *Circos* plot, Figure S8.

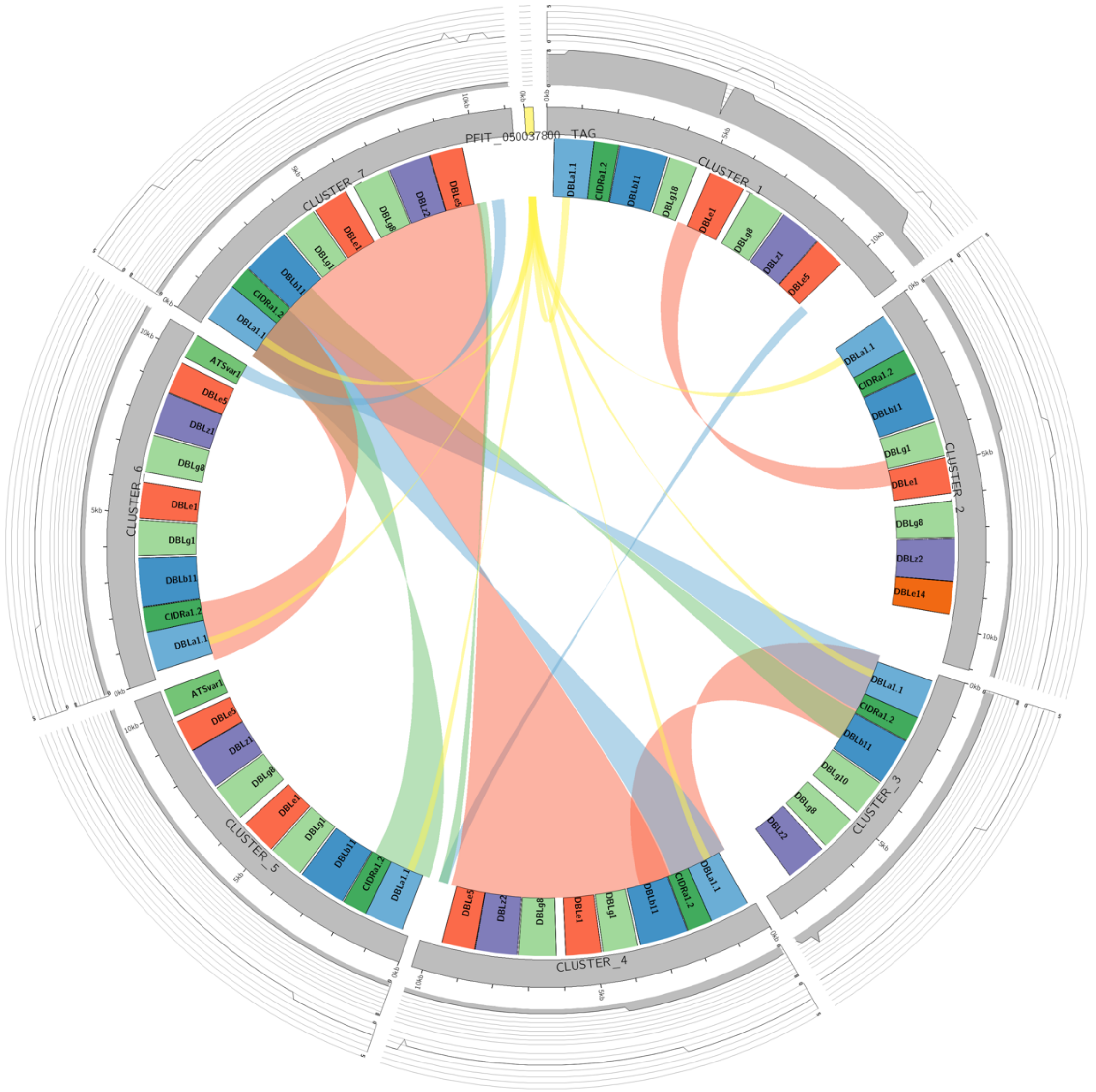

**Figure S7.** Example of Varia_VIP output showing the *Circos* plot for input tag PfIT_050037800 for the example directory.

*Example input and output of Varia_GEM*

Although the strength of the Varia_GEM module is to summarise the results of a large input set of DBLα domains, Figure S8 should the prediction of the domain configuration of the example sequence.

| | | | | | | | | | | |
|---|---|---|---|---|---|---|---|---|---|---|
| | Total reads | Singletons | | | | | | | | |
| | 1 | 1 | | | | | | | | |
| | | | | | | Position D1 | Position D1 | Position D2 | Position D2 | Position D2 |
| Sample_ID | Read_count | Blast_Cluster_ | Best Blast hit | E-value | # hits in varDB | NTSA | NTSB | DBLa0 | DBLa1 | DBLa2 |
| IT05_1_Cluste | 1 | CTTGCACGAAG | DBLa1.1 | 1.266E-128 | 62 | 54 | 2 | 4 | 55 | 0 |
| | | | | | # hits in varDB | NTSA | NTSB | DBLa0 | DBLa1 | DBLa2 |
| | | | | Sample_total: | 62 | 54 | 2 | 4 | 55 | 0 |
| Interpreded Results | | | | Position D1 | Position D1 | Position D2 | Position D2 | Position D2 | Position D2 | Position D3 |
| Sample_ID | Read_count | | | NTSA | NTSB | DBLa0 | DBLa1 | DBLa2 | DBLpam1 | CIDRa1 |
| IT05_1_Cluste | 1 | | | 1 | 0 | 0 | 1 | 0 | 0 | 1 |
| | | | Relative Expre | 1 | 0 | 0 | 1 | 0 | 0 | 1 |
| Suggested Composition | | | | | | | | | | |
| Sample_ID | Read_count | Position D1 | Position D2 | Position D3 | Position D4 | Position D5 | Position D6 | Position D7 | Position D8 | Position D9 |
| IT05_1_Cluste | 1 | NTSA1 | DBLa1.1 | CIDRa1.2 | DBLb11 | DBLg1 | DBLe1 | DBLg1 | DBLz | DBLe1 |

**Figure S7.** Example of Varia_GEM excel output of input tag PfIT_050037800. In the lower part of the figures the user can see the final prediction of the domain configuration, while on top the different counts are presented.
